# Balancing Strengths and Weaknesses in Dimensional Psychiatry

**DOI:** 10.1101/207019

**Authors:** Lindsay M. Alexander, Giovanni A. Salum, James M. Swanson, Michael P. Milham

**Affiliations:** Center for the Developing Brain, Child Mind Institute, New York, NY, USA; Universidade Federal do Rio Grande do Sul, Porto Alegre, Brazil; Child Development Center, University of California Irvine; Center for Biomedical Imaging and Neuromodulation, Nathan S. Kline Institute for Psychiatric Research, Orangeburg, NY, USA

## Abstract

**Objective:** To evaluate the feasibility and value of creating an extensible framework for psychiatric phenotyping that indexes both strengths and weaknesses of behavioral dimensions. The Extended Strengths and Weaknesses Assessment of Normal Behavior (E-SWAN) reconceptualizes each diagnostic criterion for selected DSM-5 disorders as a behavior, which can range from high (strengths) to low (weaknesses). Initial efforts have focused on Panic Disorder, Social Anxiety, Major Depression, and Disruptive Mood Dysregulation Disorder.

**Methods:** Data were collected from 523 participants (ages: 5-21 years old) in the Child Mind Institute Healthy Brain Network − an ongoing community-referred study. Parents completed each of the four E-SWAN scales and traditional unidirectional scales addressing the same disorders. Distributional properties, Item Response Theory Analysis (IRT) and Receiver Operating Characteristic (ROC) curves (for diagnostic prediction) were used to assess and compare the performance of E-SWAN and traditional scales.

**Results:** In contrast to the traditional scales, which exhibited truncated distributions, all four E-SWAN scales were found to have near-normal distributions. IRT analyses indicate the E-SWAN subscales provided reliable information about respondents throughout the population distribution; in contrast, traditional scales only provided reliable information about respondents at the high end of the distribution. Predictive value for DSM-5 diagnoses was comparable to prior scales.

**Conclusion:** E-SWAN bidirectional scales can capture the full spectrum of the population distribution for DSM disorders. The additional information provided can better inform examination of inter-individual variation in population studies, as well as facilitate the identification of factors related to resiliency in clinical samples.

## 1. INTRODUCTION

Myriad questionnaires are available for measuring psychiatric illness dimensionally. However, the vast majority are based on detection of the presence of problematic behaviors and symptoms. Although useful from a clinical perspective, the tendency to focus on only ‘one end’ of the distribution (i.e., the pathologic trait range) limits the ability of such tools to distinguish individuals from one another in less symptomatic or non-affected segments of the population (i.e., the distribution is truncated)^1,2^. This failure to consider differences in strengths among individuals is particularly problematic for psychiatric research, where efforts to model brain-behavior relationships are increasingly turning to broader community and transdiagnostic samples^3^.

The Strengths and Weaknesses Assessment of ADHD Symptoms and Normal Behavior (SWAN) provides a potentially valuable model for bidirectional questionnaire design^4^. Rather than attempting to quantify only the presence of ADHD symptoms, the SWAN probes a range of behaviors to identify relative strengths (i.e., abilities, which are indicative of adaptive behavior) and weaknesses (i.e., disabilities, which are indicative of problems requiring clinical attention, such as ADHD). This was accomplished by: 1) converting each DSM-IV ADHD symptom into a behavior and 2) expanding the typical 4-point scale of symptom presence (“not at all” to “very much”) to a 7-point scale (“far below average” to “far above average”). Numerous published studies have demonstrated that the SWAN generates bidirectional distributions that are near-normal^5–7^. Importantly, among individuals with ADHD symptomatology (i.e., a clinical sample), there is generally a high degree of agreement between the SWAN and traditional scales^1^.

Here we report on the initial design and feasibility testing of the Extended Strengths and Weaknesses Assessment of Normal Behavior (E-SWAN) — a framework that extends the general methodology of the SWAN to questionnaires for other psychiatric disorders. Consistent with the SWAN, the clinical wisdom embodied in the DSM-5 was taken as the departure point to develop each scale. Four DSM-5 disorders were chosen to provide a sampling of challenges that can arise in the conversion of DSM symptoms to dimensional probes. Major Depressive Disorder and Social Anxiety were chosen for their high prevalence in the general population^8,9^. Disruptive Mood Dysregulation Disorder (DMDD) was chosen as this new disorder in DSM-5 does not have empirically defined criteria or many valid measures for assessing symptoms^10^. Finally, Panic Disorder was chosen to determine the feasibility of applying this framework to a disorder with physiologic symptoms^11^.

The present work makes use of initial data obtained in the Child Mind Institute Healthy Brain Network sample (ages: 5-21; N=523) that enabled comparison of E-SWAN results with those obtained using equivalent unidirectional questionnaires in the same individuals. Item response theory analyses are included to demonstrate the added value of the information obtained via the E-SWAN. Additionally, we obtained informant and self-report data via the Prolific Academic platform to verify the bidirectional distributional properties of the E-SWAN in an independent sample with distinct characteristics (n=250).

## 2. METHODS

### 2.1 Questionnaire Construction: Process

The present work focused on the development and testing of four E-SWAN questionnaires (Major Depression, Disruptive Mood Dysregulation, Social Anxiety, Panic Disorder) using a uniform method based on that previously employed for construction of the SWAN (see Figure 1). *First*, each DSM-5 criterion was broken down to reflect specific symptoms that are core to each of the DSM-5 disorders. *Second*, each specific symptom was transformed into its underlying ability or behavior, i.e., the ability/behavior that when impaired or dysfunctional gives rise to the symptom. *Lastly*, each item was worded to be answered on a 7-point scale representing deviation from children of the same age, following the statement: “*When compared to children of the same age, how well does this child*…” (See Figure 1 for detailed workflow).The results from this process were discussed by a committee of experts and the final versions were circulated to experienced clinicians for comments.

**Figure 1.**
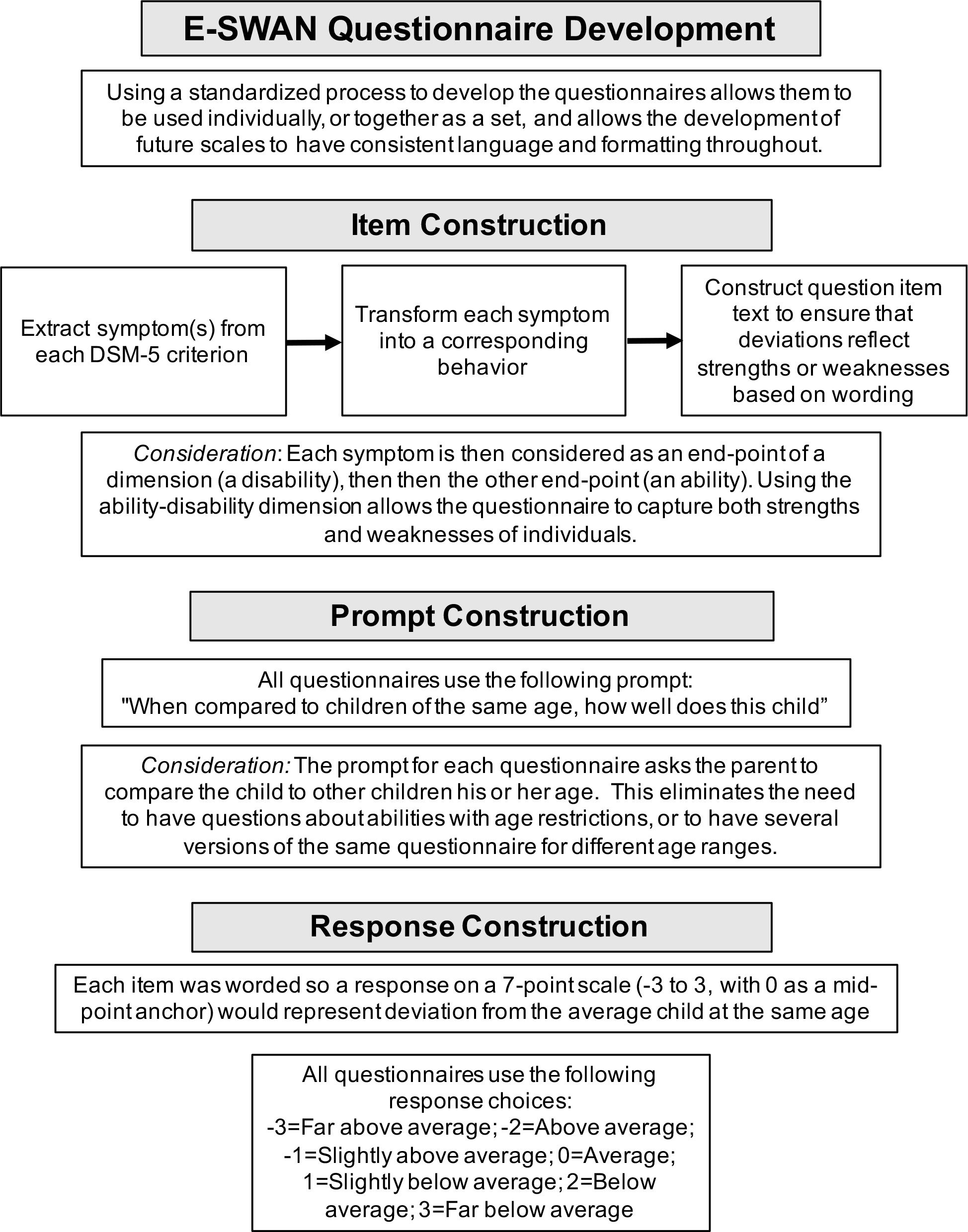
*E-SWAN Questionnaire Development Workflow*. Workflow diagram detailing the steps followed for developing the items of each E-SWAN questionnaire.

### 2.2 Questionnaire Construction: Considerations

#### 2.2.1 Level of detail and nuance

When converting DSM criteria to behaviors, we worked to ensure that question items capture the level of detail and nuance of the original criteria. This is essential as even slight changes can impact the interpretation of and responses to an item. An example of the importance of this consideration is in the DMDD questionnaire. A key criterion of a DMDD diagnosis is that the behaviors must be present in more than on setting. Initially, we indicated in each question that the behavior being rated must have taken place in more than one setting. However, we found this to be problematic, as it required parents to think about a behavior over several settings at once, and was not informative as to the specific setting(s) in which the behavior is actually taking place. As a result, we changed the questionnaire to ask each question separately for each of the settings (home, school, and with friends).

#### 2.2.2 Multiple phrases versus single

DSM diagnoses differ in number and complexity of criteria. Some DSM diagnoses have simple, clearly defined criteria, such as a symptom count, while other diagnoses contain long phrases capturing many symptoms or multiple contexts in one criterion. For example, when looking at ADHD, Depression or Social Anxiety, individual criteria are generally single symptoms. In contrast, for DMDD, each criterion encapsulates multiple symptoms. The first DMDD criterion reads “Severe recurrent temper outbursts manifested verbally and/or behaviorally that are grossly out of proportion in intensity or duration to the situation or provocation.” This one criterion captures several behaviors or symptoms across multiple settings. In the first step of the E-SWAN construction process, complex criteria such as this are broken down into multiple single items. For this one DMDD criterion, we determined that five symptoms or characteristics are being queried: verbal outbursts, behavioral outbursts, intensity, duration, and situation.

#### 2.2.3 Conditional Criteria

Some DSM diagnoses have conditional criteria that cannot easily be translated into abilities or strengths. For example, Panic Disorder criteria are mostly physiological symptoms experienced during a panic attack. To address this, we first developed three questions focused on the presence and severity of panic attacks (phrased as “moments of intense fear or discomfort”). We then ask the parent to rate how well their child is able to regulate the physiological symptoms while experiencing “a moment of intense fear or discomfort”. This allows us to potentially capture what prevents a panic attack in one individual in the same context that elicits a panic attack in another individual.

#### 2.2.4 PROMIS Guidelines

In addition to the principles that we developed, we followed the PROMIS Instrument Development Validation Scientific Standards (http://www.healthmeasures.net/images/PROMIS/PROMISStandards_Vers2.0_Final.pdf) - a set of guidelines proposed by NIH for the development of standardized assessments and rating scales. In particular, we focused on three PROMIS criteria: clarity, precision, and general applicability. Clarity means that each item is straightforward and easy to understand; not vague, confusing, or complex. To meet this goal, we used simple language characteristic of a fourth-grade reading level. Precision means that each question is specific, asking only about one behavior in one setting. We did not include multiple behaviors in one item. To meet this goal, several of the original questions were broken down into multiple questions. General applicability means that the questions do not require cultural or contextual knowledge.

### 2.3 Participants

#### 2.3.1 CMI Healthy Brain Network

Data were collected from 523 participants of the Healthy Brain Network (HBN; http://fcon_1000.projects.nitrc.org/indi/cmi_healthy_brain_network/), which is designed to be a sample of 10,000 children and adolescents from the New York City area, collected using a community-referred model that recruits based on the presence of behavioral concerns (REF). Participants selected for the present work ranged in age from 6.0-17.0. As part of the HBN protocol, parents of participants completed all E-SWAN scales (Depression, Social Anxiety, Panic Disorder, DMDD, and ADHD [SWAN]). Additionally, they completed traditionally designed instruments to assess these same disorders, including: the Mood and Feelings Questionnaire (MFQ)^12^, the Screen for Child Anxiety and Related Disorders (SCARED)^13^, the Affective Reactivity Index (ARI)^14^ and the Strengths and Difficulties Questionnaire (SDQ)^15^. DSM-5 diagnoses were established for all participants using a computerized version of the Kiddie Schedule for Affective Disorders and Schizophrenia (KSADS) (https://clinicaltrials.gov/ct2/show/NCT01866956) that was administered by licensed/license-eligible clinicians.

#### 2.3.2. Prolific Academic Sample

To confirm the generality of distributional properties for the E-SWAN, adadditional data were collected from 250 parents through Prolific Academic (an online crowdsourced data collection tool) (https://www.prolific.ac/). Users were screened based on having a child in the 6.0-17.0 age range. Parents then completed four questionnaires anonymously through a Google survey. These respondents completed the E-SWAN questionnaires only, and were given a small monetary compensation.

### 2.4 Statistical Analysis

*E-SWAN Distributional Properties*. For each of the bidirectional E-SWAN scales and corresponding traditional unidirectional scales, mean, median, skewness, and kurtosis were calculated.

*Testing Correspondence between E-SWAN and Traditional Scales*. For each of the E-SWAN scales, and its unidirectional counterpart, we calculated: 1) Cronbach’s alpha coefficients to measure internal consistency, 2) Kendall’s W coefficients to measure reliability of the different measures, and 3) Pearson correlation coefficients. To further examine the added value of measuring both strengths and weaknesses, for each questionnaire, we subdivided participants into those with mean E-SWAN scores higher for strengths (E-SWAN mean score < 0) and those with mean E-SWAN scores higher for weaknesses (E-SWAN mean score > 0), and then calculated each of these statistics again for each of the two groups (Cronbach’s alpha, Kendall’s W, Pearson correlation). Finally, we tested whether correlations between E-SWAN scales and traditional scales vary as a function of E-SWAN score using quantile regressions^16^.

*Item Response Theory (IRT)*. Next, we tested the performance of individual items from the E-SWAN and their traditional counterparts in measuring the latent trait using IRT. Prior to the IRT analysis, Confirmatory Factor Analysis (CFA) models were fitted to test the assumption of unidimensionality. These models used mean and variance adjusted weighted least squares (WLSMV) estimators, which account for the ordinal nature of the data. From this we assessed model fit using the Comparative Fit Index (CFI), the Tucker-Lewis Index (TLI) and Root Mean Square Error of Approximation (RMSEA). Values of the CFI and TLI equal to or higher than 0.90 represent an acceptable fit, and higher than 0.95 represent a good fit. Values of the RMSEA equal to or lower than 0.08 represent an acceptable fit, and lower than 0.05 represent a good fit^17,18^.

For IRT analyses, we used Graded Response Models (GRM) to calculate each item’s discrimination parameter, which indexes the strength of the relationship between each item and the latent trait, and each item difficulty parameter, which indexes in each area of the latent trait the item that concentrates the ability to provide information^19^. Test Information Function (TIF) curves were plotted for each instrument. These plots depict how well the overall test discriminates individuals, and the precision of the measurement at various levels of the latent trait. Similarly, Item Information Function (IIF) curves were plotted, which shows how each individual item on the questionnaire performs. Finally, Item Characteristic Curves (ICC) were plotted for each item of each scale. These plots depict the difficulty parameter, which represents the probability of a respondent endorsing each of the response options (e.g., far above average to far below average) along the latent trait for each instrument.

*ROC Curves for Diagnostic Prediction*. To determine the ability of E-SWAN scales to predict DSM-based diagnoses, and their comparability to traditional scales, we generated Receiver Operating Characteristic (ROC) curves for all scales using clinician consensus diagnoses from the KSADS COMP (a computerized DSM-5-based KSADS tool)^20^. We then calculated and compared Area Under the Curve (AUC) between E-SWAN scales and unidirectional scales measuring the same disorder.

All analyses were carried out in R using the following packages: lavaan^21^, ltm^22^, psych^23^, quantreg^24^, irr^25^ and pROC^26^.

## 3. RESULTS

### 3.1 Distributional Properties

Consistent with the SWAN, all E-SWAN scales were approximately normally distributed, in contrast to their traditionally designed counterparts (Figure 2 and Table 1). Kendall’s W, Cronbach’s alpha and Pearson correlation coefficients showed significant concordance, reliability, and correlation between ratings on the E-SWAN subscales and their unidirectional counterparts, respectively. To examine the added benefit of the bidirectional scales, we split the E-SWAN data into those that scored at or above 0 (higher levels of symptomatology) and those that scored below 0 (low symptomatology; higher levels of strengths). We again calculated Kendall’s W, Cronbach’s alpha, and Pearson correlations on the two groups. We found that reliability, concordance and correlations were markedly lower among those that scored below 0 (strengths group). This indicates that the unidirectional measures no longer align with the E-SWAN scales when measuring positive behaviors rather than symptoms. This relationship was consistently seen across all five domains assessed by the E-SWAN (DMDD, Depression, Social Anxiety, Panic Disorder, ADHD) (Table 1). Quantile regression showed that the traditional scales vary as a function of the E-SWAN scores. Stronger correlations are seen between the traditional scales and the E-SWAN scales at the extreme (pathologic) end of the trait (Supplementary Figure 1). A sample of reports from 250 parents on Prolific Academic yielded highly similar distributional properties, confirming that our findings were not specific to HBN (Supplementary Figure 2).

**Figure 2.**
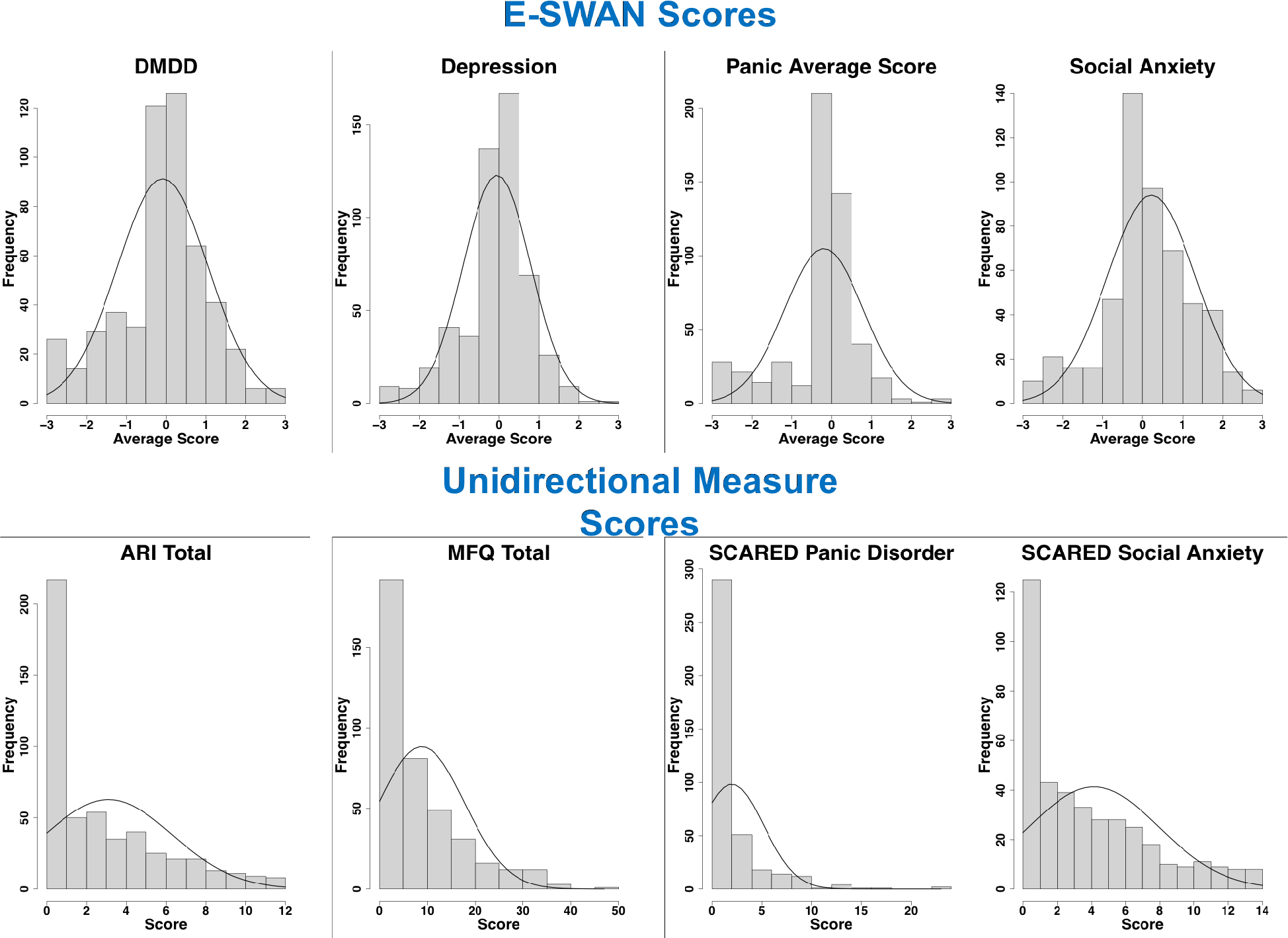
*Distribution of E-SWAN Scores and Unidirectional Measure Scores*. Distribution of scores from E-SWAN scales, shown in the top panel, compared with the distribution of scores from their unidirectional counterparts, shown in the bottom panel.

**Table 1.**
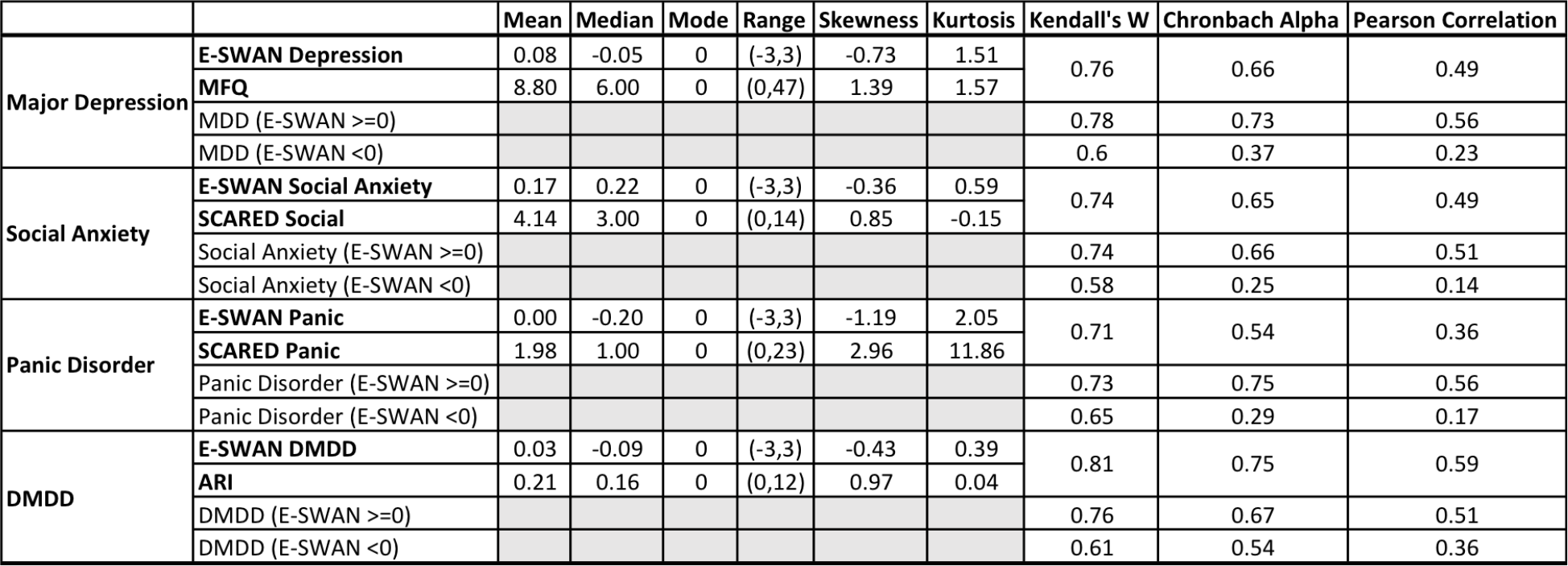
*Descriptive Statistics and Measures of Concordance and Reliability*. Measures of concordance and reliability are shown between each E-SWAN scale and its unidirectional counterparts for all participants, and for participants grouped by mean E-SWAN score.

### 3.2 Confirmatory Factor Analyses

A prerequisite for IRT analyses is the demonstration that the data being analyzed meet assumptions of sufficient unidimensionality. In this regard, scree plots were used to confirm that all E-SWAN measures, ARI, SCARED, and MFQ questionnaires met the assumption of unidimensionality (Supplementary Figure 3). CFA results are shown in Table 2. Some models showed residual correlations between items of similar content. All models showed good fit based on CFIand TLI (all values above 0.9), and most showed acceptable to good fit for RMSEA (all values <= 0.08) (Supplementary Table 1).

### 3.3 Item Response Theory Analyses

Figure 3 shows the Test Information Function curves and reliability for each scale. As can be seen in this figure, the E-SWAN scales provide reliable information across the full latent trait, from -3 to 3 standard units above the mean (reliability values 0.77-0.97 [supplementary tables 2–9]). The unidirectional scales only capture reliable information from 0 to 3 standard units above the mean of the latent trait.

**Figure 3.**
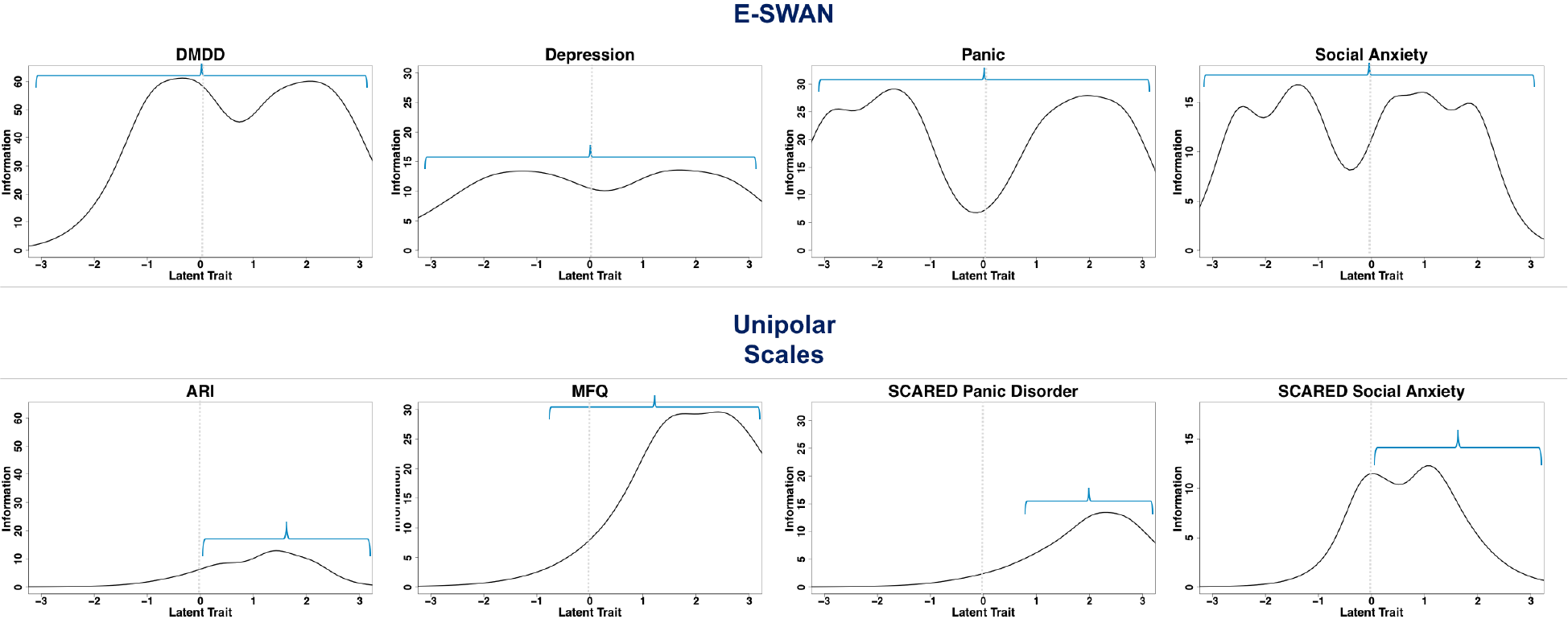
*Test Information Function Plots*. Plots depict the overall performance of each E-SWAN scale and its unidirectional counterpart at measuring the latent trait. Blue lines represent areas of the curve where information is reliable.

The Item Information Curves in Supplementary Figure 4 show that this same relationship exists for the individual items that make up the E-SWAN and unidirectional questionnaires. The E-SWAN questions capture information across the full latent trait, while the unidirectional scales only capture information at the high end of the latent trait.

Supplementary Figure 5 shows an example of the Item Characteristic Curve for one question on the E-SWAN Depression subscale and one question on the MFQ. These curves show the probability of a particular answer choice being endorsed at each level of the latent trait. Supplementary tables 2-9 indicate at which point along the latent trait there is a 50% probability of transitioning to the next response choice.

**Figure 4.**
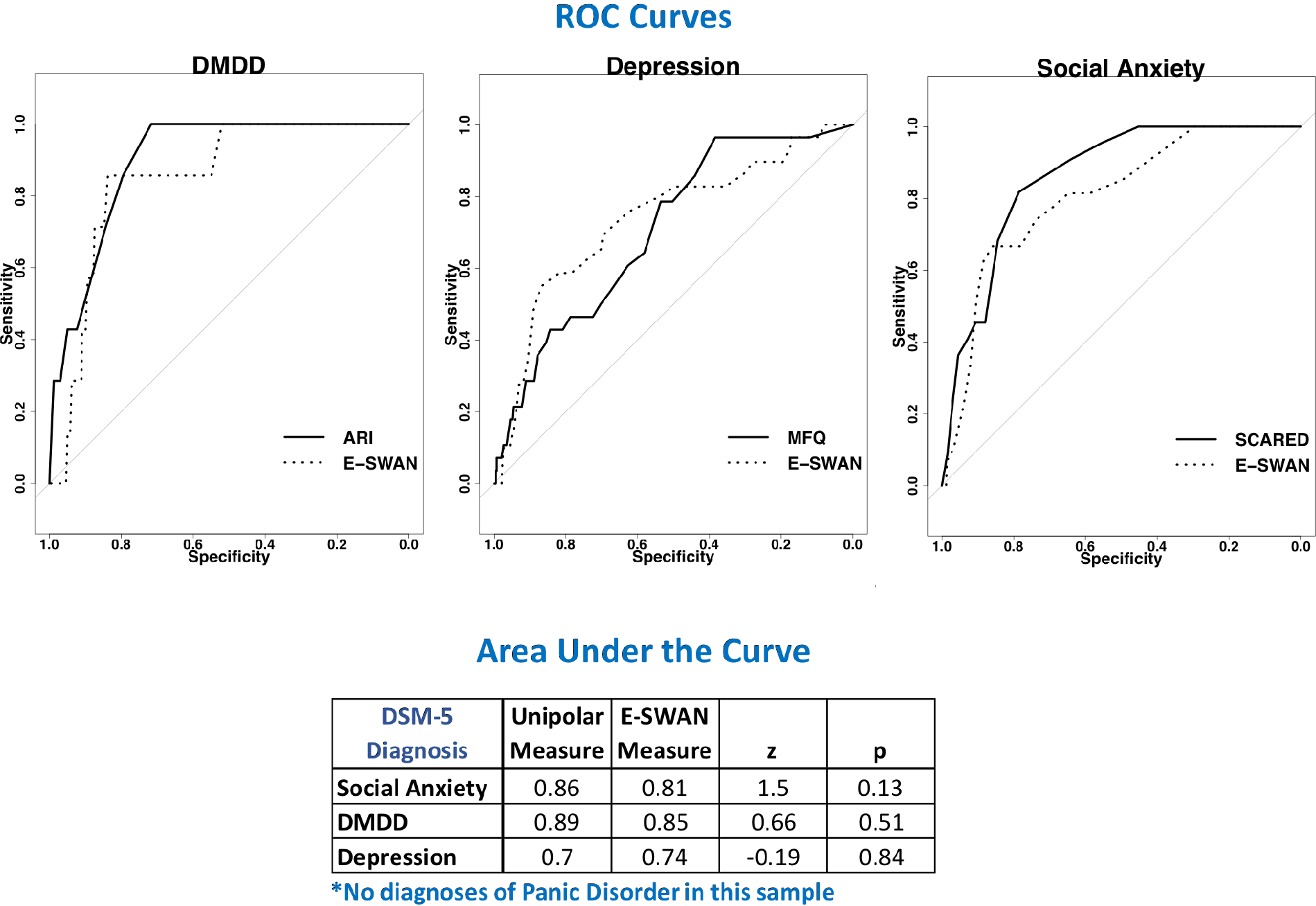
*ROC Curves*. Receiver Operating Characteristic curves representing diagnostic capabilities of both E-SWAN and unidirectional scales for DMDD, Depression, and Social Anxiety.

### 3.4 Predictive Value for DSM Diagnosis

A key concern that may arise regarding the E-SWAN is whether the resulting scales have comparable predictive value for DSM diagnoses relative to previously established unidirectional scales. Given the high correlation of scores at the high end of the latent trait, one would expect this to be the case. To test this, we generated ROC curves for all scales using diagnoses generated from the K-SADS (Figure 4). Both E-SWAN and traditional scales performed well (AUC values 0.7-0.89), indicating that they are comparable screening tools and giving increased support for the validity of the E-SWAN questionnaires.

## 4. DISCUSSION

Inspired by the SWAN, we developed and tested a generalized framework for constructing questionnaires to assess the full range of behavior defined by DSM symptoms, when considered as an endpoint of a dimension. Consistent with the SWAN, each of the E-SWAN questionnaires were constructed to be bidirectional, i.e., indexing both strengths (abilities) and weaknesses (disabilities). When compared to the unidirectional scales, the E-SWAN scales exhibited distributional properties that were near-normal rather than highly skewed or truncated. As predicted, for each trait, a strong correspondence was noted between the E-SWAN scores and traditional scale scores among individuals at the high (pathological) end, but not at the low end. IRT analyses suggested that in contrast to traditional scales, the E-SWAN subscales exhibited good discrimination and reliability across the full latent trait (z-scores from −3 to +3; reliabilities ranging from 0.77 to 0.97) — not just at the high end. Finally, we demonstrated the ability to generate self-report questionnaires using the E-SWAN framework. Consistent with the data from Healthy Brain Network participants, our online sample from Prolific Academic yielded a near-normal distribution, although shifted slightly to the left (i.e., less symptomatic), as would be expected given the differences in recruitment strategies (online crowdsourcing community vs. community-referred based on the presence of behavioral concerns).

The ability to meaningfully and reliably catalog variance across the entirety of a population becomes more important as biological and epidemiologic studies shift away from categorical, syndromic characterizations of psychiatric illness. Efforts such as the NIMH Research Domain Criteria Project have successfully drawn attention to the potential added value of dimensional characterizations that cut across the diagnostic boundaries specified by DSM and ICD^3^. Yet the vast majority of questionnaires focused on mental health are limited in their ability to characterize variation among individuals beyond the symptomatic segment of the population^1,2^. As demonstrated in the present work, the E-SWAN framework offers a viable alternative for improving our ability to differentiate individuals that are non-symptomatic in a given domain. It does so without losing track of the clinical significance of identifying a pathological range of the trait (which is used as the departure point for characterizing the trait). The data from the E-SWAN are more statistically appropriate for dimensional analysis. While the present work used the total score obtained for a given scale as the unit of analysis, future work may benefit from consideration of individual item scores. Similar to overall questionnaire scores, individual items of the E-SWAN are intended to represent a bidirectional dimension. This property can be particularly valuable for efforts focused on the identification of abilities that may have protective effects or confer resilience, as well as disabilities associated with impairment. For example, when evaluating individuals with equivalent symptom profiles (defined by one end of the underlying bidirectional dimensions), though differing outcomes, the presence or absence of strengths (on the other end of the dimensions — but not measured by traditional scales) may result in different outcomes. The E-SWAN scales can be used to capture such distinctions.

There are a number of limitations of the E-SWAN framework that suggest areas for improvement. First, a key assumption of the E-SWAN framework — that the underlying dimension of behavior described is bidirectional and normally distributed in the general population — may not hold for all disorders. While likely reasonable for most DSM disorders, some, such as PTSD and Substance Use Disorders, represent clear instances where this is not the case, as only a subset of the population has had exposure to trauma or a given substance^11^. For these disorders, the prompts and range of responses can be changed to create a distribution in a subset of the population defined by the presence of a particular exposure (e.g., stressor, substance use). Questionnaires for both of these disorders are under development (see eswan.org for current drafts). Second is the potential for biased reporting, which can arise from either a skewed perception of ‘other children the same age’ on the part of the informant, or a bias to see a child as more (or less) able than they are (e.g., the “Lake Wobegon Effect”)^27^. Arguably such biases are also present in unidirectional questionnaires, though centered more around ratings of frequency. As demonstrated with other questionnaire tools, one of the most promising ways of overcoming such bias is the collection of data from multiple informants.

As demonstrated in our comparison of results from Healthy Brain Network and Prolific Academic, the mean of the distribution obtained in a given sample can vary depending on the specific segment of the population sampled. Within a given study, this is not necessarily a problem. However, if not taken into account, such variation can lead to confounds or biases when attempting to compare or combine across studies, or to develop clinical cutoffs. An effective solution would arguably be the generation of appropriately-sized normative samples to serve as a reference. This need is not different from any other questionnaire. A positive aspect of the E-SWAN versus other questionnaires is that one can more easily compare distributional characteristics to understand differences among samples in their entirety.

In the spirit of collaboration and open science, all E-SWAN questionnaires are freely available for use and can be accessed at www.eswan.org, licensed Creative Commons (CC) BY 4.0 to encourage maximal dissemination and application of the questionnaires. It is our hope that other investigative teams will join in the effort to create the full range of E-SWAN questionnaires encompassing all major psychiatric syndromes.

## Supplemental Figure Legends

**Supplementary Figure 1.**
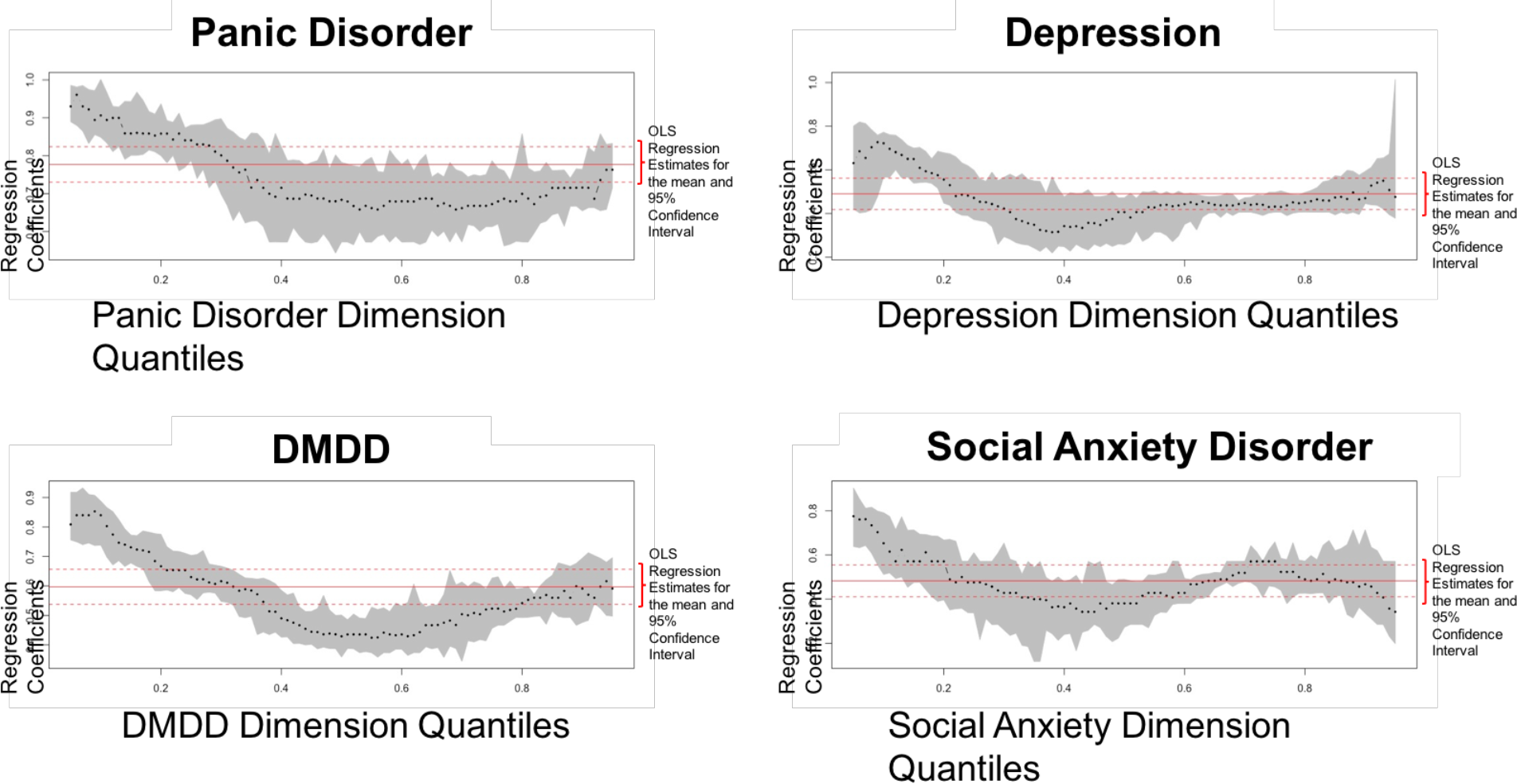
*Quantile Regression Plots*. Quantile regression plots showing that the traditional scales vary as a function of the E-SWAN scores. Stronger correlations are seen between the traditional scales and the E-SWAN scales at the extreme (pathologic) end of the trait.

**Supplementary Figure 2.**
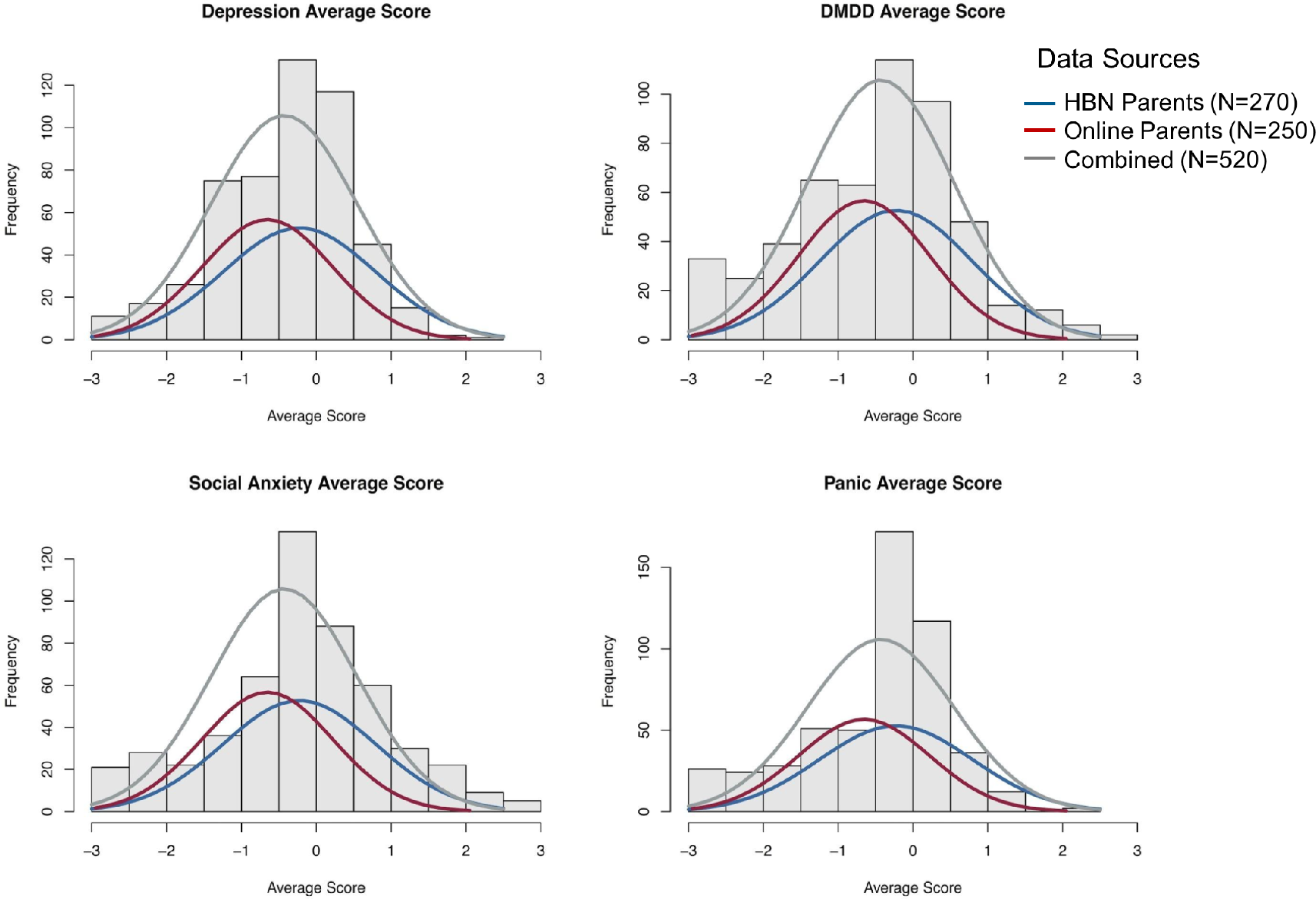
*Prolific Academic Parent Report and HBN Parent Report*. Plots comparing distribution of E-SWAN scores from HBN parent reports and online parent reports gathered through Prolific Academic. Both samples show similar distributions, with the online sample having a slightly lower mean.

Supplementary Figure 3. *Scree plots*. Scree plots indicate that all scales meet the assumption of unidimensionality required for CFA analyses.

**Supplementary Figure 4.**
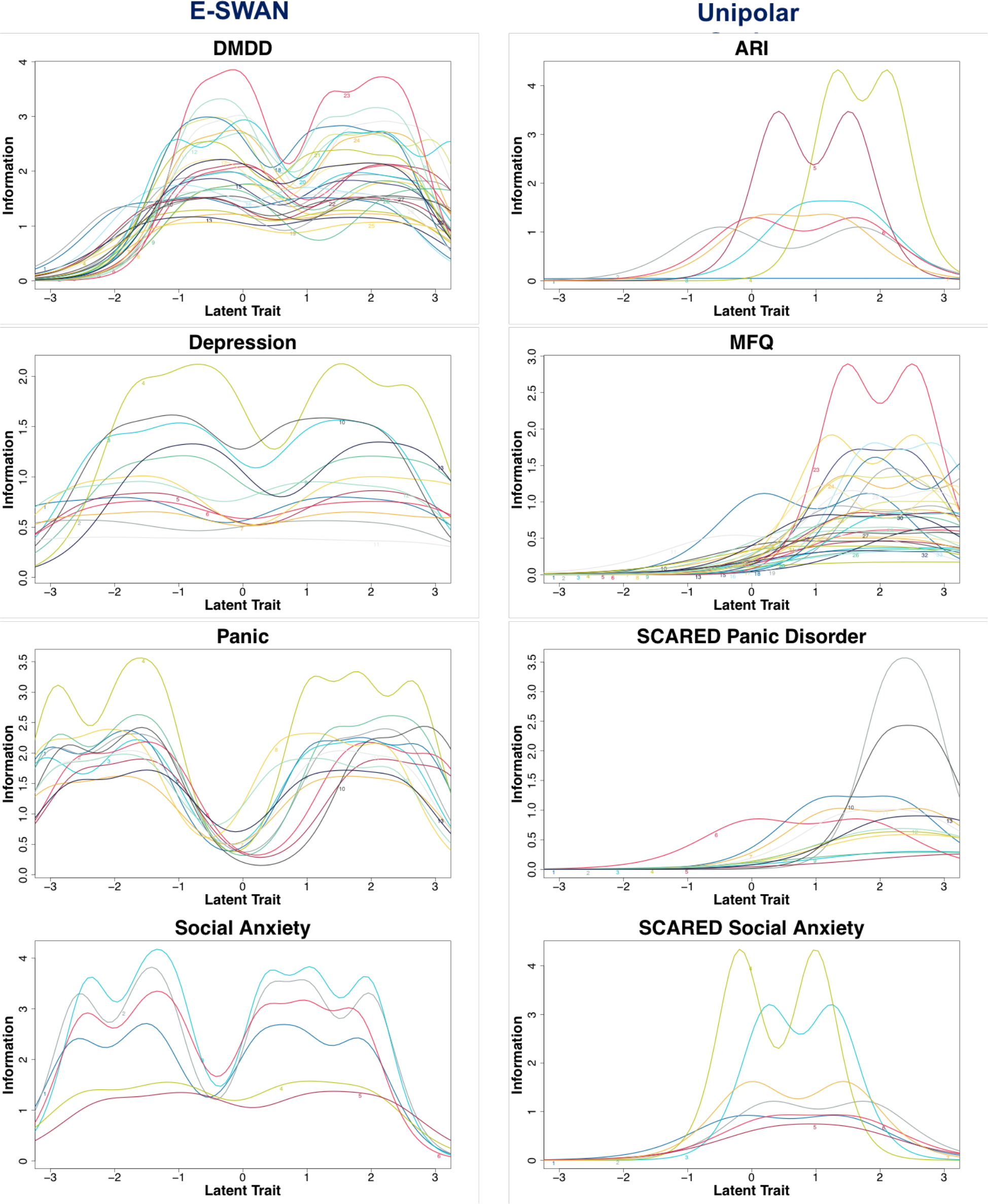
*Item Information Curves*. Plots depict the performance of each item on each of the E-SWAN scales and its unidirectional counterpart at measuring the latent trait.

**Supplementary Figure 5.**
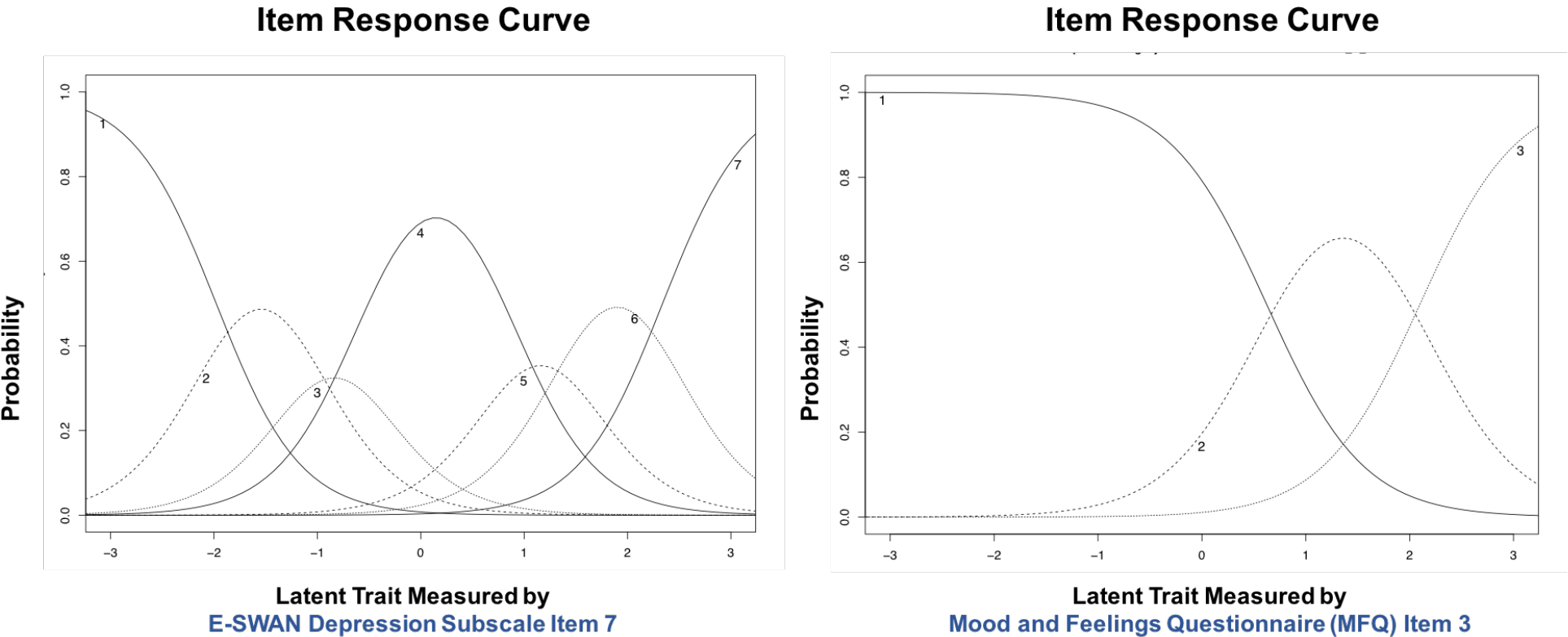
*Sample Item Characteristic Curves*. An example Item Characteristic Curve from the E-SWAN Depression scale, and the MFQ are shown. Lines represent the probability of endorsing a specific response choice at each area of the latent trait. For the sample item from the E-SWAN (item 7 of E-SWAN Depression scale) there is a 50% estimated probability that a participant will transition from an answer of ‘Far Below Average’ to ‘Below Average’ or higher in this item when latent trait levels are 2.06 standard units below the mean, 50% probability of transitioning from ‘Below Average’ to ‘Slightly Above Average’ when latent trait levels are 1.24 standard units below the mean, and 50% probability of transitioning from ‘Slightly Below Average’ to ‘About Average’ when latent trait levels are 0.7 standard units below the mean. Categories of ‘Slightly Above Average’ or higher, ‘Above Average’ or higher and ‘Far Above Average’ are endorsed with a 50% probability when latent trait levels are 0.6, 1.37, and 2.18 standard units above the mean, respectively. For the sample item of the MFQ (item 3 of the MFQ), there is a 50% estimated probability that a participant will transition from an answer of ‘Not True’ to ‘Sometimes’ or higher when latent trait levels are 1.58 standard units above the mean, and 50% probability of transitioning from ‘Sometimes’ to ‘True’ when latent trait levels are 3.49 standard units above the mean.

**Supplementary Table Legends:**

**Supplementary Table 1.** *Measures of Fit for Confirmatory Factor Analysis*.

|  | Comparative<br>Fit Index (CFI) | Tucker-Lewis<br>Index (TLI) | RMSEA |
| --- | --- | --- | --- |
| **E-SWAN Depression** | 0.99** | 0.99** | 0.085<sup>^</sup> |
| **E-SWAN Social Anxiety** | 1.00** | 1.00** | 0.014** |
| **E-SWAN Panic Disorder** | 0.99** | 0.99** | 0.082<sup>^</sup> |
| **E-SWAN DMDD** | 0.99** | 0.99** | 0.059** |
| **ARI** | 1.00** | 0.99** | 0.035** |
| **MFQ** | 0.98** | 0.98** | 0.056** |
| **SCARED Panic** | 0.99** | 0.99** | 0.022** |
| **SCARED Social** | 0.99** | 0.99** | 0.037** |

\*acceptable; \*\*good

**Supplementary Table 2.** *Item Response Theory Parameters for E-SWAN Depression Scale*.

| | Discrimination ($\alpha$) and difficulty ($\beta$) parameters | | | | | | | | Item information by area of the latent trait (z-score) | | | | | | |
| --- | --- | --- | --- | --- | --- | --- | --- | --- | --- | --- | --- | --- | --- | --- | --- |
| | $\alpha$ | $\beta_1$ | $\beta_2$ | $\beta_3$ | $\beta_4$ | $\beta_5$ | $\beta_6$ | Mean($\beta$) | -3 | -2 | -1 | 0 | 1 | 2 | 3 |
| MDD_1A | 1.612 | -3.237 | -2.036 | -1.364 | 0.872 | 1.534 | 2.614 | -0.270 | 0.37 | 0.48 | 0.54 | 0.55 | 0.52 | 0.44 | 0.32 |
| MDD_1B | 1.35 | -3.488 | -2.546 | -1.771 | 0.344 | 1.33 | 2.45 | -0.614 | 0.34 | 0.42 | 0.47 | 0.47 | 0.42 | 0.35 | 0.25 |
| MDD_2A | 2.251 | -2.185 | -1.207 | -0.579 | 0.984 | 1.564 | 2.282 | 0.143 | 0.51 | 0.77 | 0.86 | 0.83 | 0.86 | 0.83 | 0.62 |
| MDD_2B | 2.654 | -1.705 | -0.884 | -0.283 | 1.254 | 1.71 | 2.666 | 0.460 | 0.48 | 0.83 | 0.97 | 0.86 | 0.83 | 0.95 | 0.87 |
| MDD_3A | 1.661 | -2.381 | -1.346 | -0.963 | 1.408 | 2.066 | 2.742 | 0.254 | 0.33 | 0.46 | 0.55 | 0.58 | 0.57 | 0.51 | 0.4 |
| MDD_3B | 1.566 | -2.431 | -1.462 | -0.93 | 1.075 | 2.052 | 2.971 | 0.213 | 0.34 | 0.47 | 0.56 | 0.59 | 0.57 | 0.5 | 0.38 |
| MDD_4 | 1.467 | -2.91 | -1.541 | -1.006 | 1.138 | 2.132 | 3.744 | 0.260 | 0.3 | 0.39 | 0.45 | 0.47 | 0.46 | 0.4 | 0.32 |
| MDD_5 | 1.822 | -2.631 | -1.581 | -1.106 | 1.353 | 2.234 | 3.303 | 0.262 | 0.36 | 0.48 | 0.54 | 0.54 | 0.53 | 0.5 | 0.41 |
| MDD_6 | 1.988 | -1.908 | -0.979 | -0.5 | 1.333 | 2.105 | 2.971 | 0.504 | 0.35 | 0.56 | 0.69 | 0.72 | 0.71 | 0.68 | 0.56 |
| MDD_7 | 2.293 | -2.064 | -1.24 | -0.712 | 0.626 | 1.37 | 2.185 | 0.028 | 0.54 | 0.82 | 0.92 | 0.88 | 0.86 | 0.77 | 0.56 |
| MDD_8A | 1.125 | -3.354 | -2.37 | -1.788 | -0.378 | 0.96 | 2.779 | -0.692 | 0.3 | 0.37 | 0.41 | 0.41 | 0.36 | 0.3 | 0.22 |
| MDD_8B | 1.766 | -2.817 | -1.719 | -1.164 | 0.577 | 1.373 | 2.487 | -0.211 | 0.46 | 0.62 | 0.7 | 0.7 | 0.66 | 0.55 | 0.39 |
| MDD_9 | 2.083 | -1.474 | -0.681 | -0.369 | 1.558 | 2.149 | 2.862 | 0.674 | 0.32 | 0.56 | 0.75 | 0.79 | 0.77 | 0.77 | 0.66 |
| Mean | 1.818 | -2.507 | -1.507 | -0.964 | 0.934 | 1.737 | 2.774 | 0.078 |  |  |  |  |  |  |  |
| Test information |  |  |  |  |  |  |  |  | 4.99 | 7.23 | 8.4 | 8.39 | 8.12 | 7.55 | 5.98 |
| SEM |  |  |  |  |  |  |  |  | 0.45 | 0.37 | 0.34 | 0.35 | 0.35 | 0.36 | 0.41 |
| Reliability |  |  |  |  |  |  |  |  | 0.8 | 0.86 | 0.88 | 0.88 | 0.88 | 0.87 | 0.83 |

**Supplementary Table 3.**
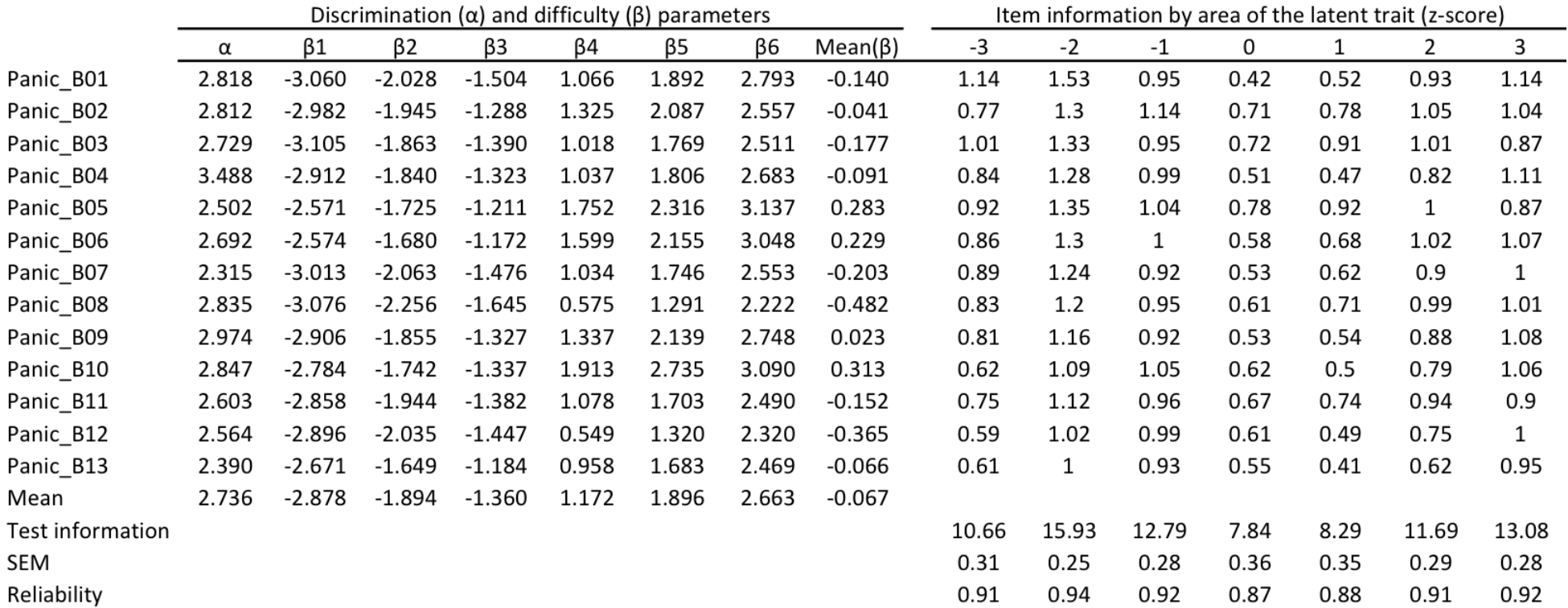
*Item Response Theory Parameters for E-SWAN Panic Disorder Scale*.

| | Discrimination ($\alpha$) and difficulty ($\beta$) parameters | | | | | | | | Item information by area of the latent trait (z-score) | | | | | | |
| --- | --- | --- | --- | --- | --- | --- | --- | --- | --- | --- | --- | --- | --- | --- | --- |
| | $\alpha$ | $\beta_1$ | $\beta_2$ | $\beta_3$ | $\beta_4$ | $\beta_5$ | $\beta_6$ | Mean($\beta$) | -3 | -2 | -1 | 0 | 1 | 2 | 3 |
| Panic_B01 | 2.818 | -3.060 | -2.028 | -1.504 | 1.066 | 1.892 | 2.793 | -0.140 | 1.14 | 1.53 | 0.95 | 0.42 | 0.52 | 0.93 | 1.14 |
| Panic_B02 | 2.812 | -2.982 | -1.945 | -1.288 | 1.325 | 2.087 | 2.557 | -0.041 | 0.77 | 1.3 | 1.14 | 0.71 | 0.78 | 1.05 | 1.04 |
| Panic_B03 | 2.729 | -3.105 | -1.863 | -1.390 | 1.018 | 1.769 | 2.511 | -0.177 | 1.01 | 1.33 | 0.95 | 0.72 | 0.91 | 1.01 | 0.87 |
| Panic_B04 | 3.488 | -2.912 | -1.840 | -1.323 | 1.037 | 1.806 | 2.683 | -0.091 | 0.84 | 1.28 | 0.99 | 0.51 | 0.47 | 0.82 | 1.11 |
| Panic_B05 | 2.502 | -2.571 | -1.725 | -1.211 | 1.752 | 2.316 | 3.137 | 0.283 | 0.92 | 1.35 | 1.04 | 0.78 | 0.92 | 1 | 0.87 |
| Panic_B06 | 2.692 | -2.574 | -1.680 | -1.172 | 1.599 | 2.155 | 3.048 | 0.229 | 0.86 | 1.3 | 1 | 0.58 | 0.68 | 1.02 | 1.07 |
| Panic_B07 | 2.315 | -3.013 | -2.063 | -1.476 | 1.034 | 1.746 | 2.553 | -0.203 | 0.89 | 1.24 | 0.92 | 0.53 | 0.62 | 0.9 | 1 |
| Panic_B08 | 2.835 | -3.076 | -2.256 | -1.645 | 0.575 | 1.291 | 2.222 | -0.482 | 0.83 | 1.2 | 0.95 | 0.61 | 0.71 | 0.99 | 1.01 |
| Panic_B09 | 2.974 | -2.906 | -1.855 | -1.327 | 1.337 | 2.139 | 2.748 | 0.023 | 0.81 | 1.16 | 0.92 | 0.53 | 0.54 | 0.88 | 1.08 |
| Panic_B10 | 2.847 | -2.784 | -1.742 | -1.337 | 1.913 | 2.735 | 3.090 | 0.313 | 0.62 | 1.09 | 1.05 | 0.62 | 0.5 | 0.79 | 1.06 |
| Panic_B11 | 2.603 | -2.858 | -1.944 | -1.382 | 1.078 | 1.703 | 2.490 | -0.152 | 0.75 | 1.12 | 0.96 | 0.67 | 0.74 | 0.94 | 0.9 |
| Panic_B12 | 2.564 | -2.896 | -2.035 | -1.447 | 0.549 | 1.320 | 2.320 | -0.365 | 0.59 | 1.02 | 0.99 | 0.61 | 0.49 | 0.75 | 1 |
| Panic_B13 | 2.390 | -2.671 | -1.649 | -1.184 | 0.958 | 1.683 | 2.469 | -0.066 | 0.61 | 1 | 0.93 | 0.55 | 0.41 | 0.62 | 0.95 |
| Mean | 2.736 | -2.878 | -1.894 | -1.360 | 1.172 | 1.896 | 2.663 | -0.067 |  |  |  |  |  |  |  |
| Test information |  |  |  |  |  |  |  |  | 10.66 | 15.93 | 12.79 | 7.84 | 8.29 | 11.69 | 13.08 |
| SEM |  |  |  |  |  |  |  |  | 0.31 | 0.25 | 0.28 | 0.36 | 0.35 | 0.29 | 0.28 |
| Reliability |  |  |  |  |  |  |  |  | 0.91 | 0.94 | 0.92 | 0.87 | 0.88 | 0.91 | 0.92 |

**Supplementary Table 4.** *Item Response Theory Parameters for E-SWAN Social Anxiety Scale*.

| | Discrimination ($\alpha$) and difficulty ($\beta$) parameters | | | | | | | | Item information by area of the latent trait (z-score) | | | | | | |
| --- | --- | --- | --- | --- | --- | --- | --- | --- | --- | --- | --- | --- | --- | --- | --- |
| | $\alpha$ | $\beta_1$ | $\beta_2$ | $\beta_3$ | $\beta_4$ | $\beta_5$ | $\beta_6$ | Mean($\beta$) | -3 | -2 | -1 | 0 | 1 | 2 | 3 |
| SocAnx_01 | 3.035 | -2.621 | -1.618 | -1.340 | 0.266 | 0.888 | 1.863 | -0.427 | 1.4 | 1.45 | 0.76 | 0.76 | 1.12 | 1.15 | 0.99 |
| SocAnx_02 | 3.584 | -2.569 | -1.578 | -1.239 | 0.377 | 1.036 | 1.998 | -0.329 | 1.3 | 1.41 | 0.92 | 0.92 | 1.17 | 1.13 | 0.93 |
| SocAnx_03 | 3.746 | -2.412 | -1.512 | -1.108 | 0.370 | 1.086 | 1.958 | -0.270 | 1.15 | 1.26 | 0.87 | 0.81 | 1 | 0.95 | 0.77 |
| SocAnx_04A | 2.263 | -2.474 | -1.397 | -0.845 | 0.582 | 1.162 | 1.948 | -0.171 | 1.03 | 1.17 | 0.94 | 0.91 | 0.99 | 0.87 | 0.61 |
| SocAnx_04B | 2.104 | -2.195 | -1.161 | -0.601 | 0.898 | 1.466 | 2.204 | 0.102 | 0.6 | 0.87 | 0.96 | 0.94 | 0.95 | 0.82 | 0.53 |
| SocAnx_05 | 3.349 | -2.480 | -1.525 | -1.095 | 0.291 | 0.991 | 1.775 | -0.341 | 0.48 | 0.77 | 0.92 | 0.93 | 0.94 | 0.86 | 0.59 |
| Mean | 3.014 | -2.459 | -1.465 | -1.038 | 0.464 | 1.105 | 1.958 | -0.239 |  |  |  |  |  |  |  |
| Test information |  |  |  |  |  |  |  |  | 5.96 | 6.93 | 5.37 | 5.27 | 6.17 | 5.78 | 4.43 |
| SEM |  |  |  |  |  |  |  |  | 0.41 | 0.38 | 0.43 | 0.44 | 0.4 | 0.42 | 0.48 |
| Reliability |  |  |  |  |  |  |  |  | 0.83 | 0.86 | 0.81 | 0.81 | 0.84 | 0.83 | 0.77 |

**Supplementary Table 5.** *Item Response Theory Parameters for E-SWAN DMDD Scale*.

| | Discrimination ($\alpha$) and difficulty ($\beta$) parameters | | | | | | | | Item information by area of the latent trait (z-score) | | | | | | |
| --- | --- | --- | --- | --- | --- | --- | --- | --- | --- | --- | --- | --- | --- | --- | --- |
| | $\alpha$ | $\beta_1$ | $\beta_2$ | $\beta_3$ | $\beta_4$ | $\beta_5$ | $\beta_6$ | Mean($\beta$) | -3 | -2 | -1 | 0 | 1 | 2 | 3 |
| DMDD_1A | 2.234 | -1.868 | -0.927 | -0.512 | 0.660 | 1.360 | 2.189 | 0.150 | 0.69 | 1.43 | 1.55 | 0.97 | 0.94 | 1.26 | 1.25 |
| DMDD_1B | 2.561 | -1.166 | -0.357 | 0.049 | 1.480 | 2.197 | 3.103 | 0.884 | 0.77 | 1.47 | 1.52 | 0.97 | 1.04 | 1.34 | 1.28 |
| DMDD_1C | 2.547 | -0.969 | -0.233 | 0.104 | 1.394 | 1.949 | 2.721 | 0.828 | 0.81 | 1.52 | 1.47 | 0.82 | 0.85 | 1.19 | 1.29 |
| DMDD_2A | 2.041 | -1.420 | -0.629 | -0.187 | 1.145 | 1.764 | 2.582 | 0.543 | 0.61 | 1.31 | 1.56 | 1.1 | 1.08 | 1.36 | 1.24 |
| DMDD_2B | 2.641 | -0.836 | -0.111 | 0.283 | 1.755 | 2.325 | 3.009 | 1.071 | 0.97 | 1.39 | 1.18 | 0.89 | 1.07 | 1.16 | 0.97 |
| DMDD_2C | 2.608 | -0.766 | -0.035 | 0.349 | 1.685 | 2.266 | 2.770 | 1.045 | 1 | 1.42 | 1.14 | 0.85 | 0.98 | 1.04 | 0.93 |
| DMDD_3A | 1.987 | -1.032 | -0.216 | 0.070 | 1.235 | 1.916 | 2.648 | 0.770 | 0.99 | 1.38 | 1.1 | 0.79 | 0.94 | 1.01 | 0.9 |
| DMDD_3B | 2.407 | -0.710 | 0.063 | 0.353 | 1.863 | 2.332 | 2.858 | 1.127 | 0.72 | 1.31 | 1.28 | 0.9 | 0.95 | 1.12 | 1.03 |
| DMDD_3C | 2.416 | -0.715 | 0.093 | 0.384 | 1.799 | 2.211 | 2.671 | 1.074 | 0.48 | 1.04 | 1.32 | 1.05 | 0.94 | 1.16 | 1.12 |
| DMDD_4A | 2.213 | -1.413 | -0.659 | -0.252 | 1.139 | 1.818 | 2.612 | 0.541 | 0.57 | 1.14 | 1.35 | 1.11 | 1.14 | 1.2 | 0.93 |
| DMDD_4B | 3.144 | -0.761 | -0.136 | 0.235 | 1.582 | 2.220 | 2.869 | 1.002 | 0.71 | 1.22 | 1.27 | 1 | 1.04 | 1.08 | 0.9 |
| DMDD_4C | 2.977 | -0.803 | -0.104 | 0.211 | 1.346 | 1.887 | 2.473 | 0.835 | 0.4 | 0.91 | 1.25 | 1.13 | 1.04 | 1.2 | 1.08 |
| DMDD_5A | 1.931 | -1.534 | -0.819 | -0.301 | 1.040 | 1.754 | 2.499 | 0.440 | 0.79 | 1.22 | 1.16 | 0.97 | 1.02 | 0.99 | 0.76 |
| DMDD_5B | 2.666 | -1.004 | -0.476 | -0.089 | 1.261 | 1.907 | 2.567 | 0.694 | 0.62 | 1.11 | 1.22 | 1.06 | 1.07 | 1.05 | 0.77 |
| DMDD_5C | 2.456 | -1.106 | -0.461 | -0.104 | 1.237 | 1.847 | 2.641 | 0.676 | 0.32 | 0.74 | 1.11 | 1.1 | 1.01 | 1.11 | 0.99 |
| DMDD_6A | 2.365 | -1.582 | -0.976 | -0.541 | 0.644 | 1.240 | 2.127 | 0.152 | 0.31 | 0.7 | 1.07 | 1.12 | 1.06 | 1.16 | 0.99 |
| DMDD_6B | 3.117 | -1.037 | -0.471 | -0.052 | 1.301 | 2.081 | 2.995 | 0.803 | 0.59 | 0.97 | 1.05 | 0.92 | 0.96 | 0.95 | 0.7 |
| DMDD_6C | 3.106 | -1.031 | -0.570 | -0.128 | 1.008 | 1.641 | 2.355 | 0.546 | 0.51 | 0.85 | 1 | 0.94 | 0.95 | 0.96 | 0.75 |
| DMDD_7A | 2.250 | -1.972 | -0.772 | -0.254 | 1.556 | 2.263 | 3.127 | 0.658 | 0.57 | 0.94 | 1.05 | 0.97 | 0.98 | 0.9 | 0.62 |
| DMDD_7B | 3.129 | -1.105 | -0.166 | 0.253 | 1.506 | 2.264 | 3.297 | 1.008 | 0.46 | 0.81 | 0.96 | 0.84 | 0.81 | 0.92 | 0.85 |
| DMDD_7C | 2.862 | -1.076 | -0.537 | -0.190 | 0.993 | 1.699 | 2.493 | 0.564 | 0.6 | 0.99 | 1.14 | 1.09 | 0.99 | 0.76 | 0.43 |
| DMDD_8A | 2.232 | -1.458 | -0.605 | -0.239 | 1.280 | 1.920 | 2.713 | 0.602 | 0.6 | 0.96 | 1.1 | 1.06 | 0.96 | 0.72 | 0.41 |
| DMDD_8B | 3.548 | -0.748 | -0.211 | 0.104 | 1.309 | 1.998 | 2.504 | 0.826 | 0.57 | 0.91 | 1.04 | 1 | 0.92 | 0.73 | 0.45 |
| DMDD_8C | 3.000 | -0.855 | -0.199 | 0.158 | 1.480 | 2.137 | 2.691 | 0.902 | 0.68 | 0.95 | 0.98 | 0.92 | 0.84 | 0.66 | 0.4 |
| DMDD_9A | 1.854 | -1.389 | -0.574 | -0.056 | 1.498 | 2.208 | 2.952 | 0.773 | 0.51 | 0.85 | 1.02 | 1.01 | 0.95 | 0.77 | 0.47 |
| DMDD_9B | 2.336 | -1.052 | -0.301 | 0.134 | 2.226 | 2.895 | 3.506 | 1.235 | 0.64 | 0.93 | 1.03 | 0.98 | 0.86 | 0.63 | 0.36 |
| DMDD_9C | 2.246 | -1.183 | -0.327 | 0.160 | 1.690 | 2.321 | 3.056 | 0.953 | 0.68 | 0.95 | 1.01 | 0.95 | 0.84 | 0.62 | 0.35 |
| DMDD_10A | 2.293 | -1.344 | -0.661 | -0.283 | 0.937 | 1.622 | 2.380 | 0.442 | 0.37 | 0.69 | 0.96 | 1.03 | 0.97 | 0.8 | 0.51 |
| DMDD_10B | 3.269 | -0.798 | -0.354 | 0.002 | 1.238 | 1.933 | 2.495 | 0.753 | 0.4 | 0.63 | 0.78 | 0.82 | 0.79 | 0.68 | 0.47 |
| DMDD_10C | 2.670 | -0.947 | -0.363 | 0.048 | 1.158 | 1.870 | 2.495 | 0.710 | 0.48 | 0.7 | 0.83 | 0.84 | 0.77 | 0.6 | 0.37 |
| Mean | 2.570 | -1.123 | -0.403 | -0.010 | 1.348 | 1.995 | 2.713 | 0.753 |  |  |  |  |  |  |  |
| Test information |  |  |  |  |  |  |  |  | 18.41 | 31.48 | 34.49 | 29.21 | 28.74 | 29.14 | 23.55 |
| SEM |  |  |  |  |  |  |  |  | 0.23 | 0.18 | 0.17 | 0.19 | 0.19 | 0.19 | 0.21 |
| Reliability |  |  |  |  |  |  |  |  | 0.95 | 0.97 | 0.97 | 0.97 | 0.97 | 0.97 | 0.96 |

**Supplementary Table 6.** *Item Response Theory Parameters for ARI*.

| | Discrimination ($\alpha$) and difficulty ($\beta$) parameters | | | | Item information by area of the latent trait (z-score) | | | | | | |
| --- | --- | --- | --- | --- | --- | --- | --- | --- | --- | --- | --- |
| | $\alpha$ | $\beta_1$ | $\beta_2$ | Mean($\beta$) | -3 | -2 | -1 | 0 | 1 | 2 | 3 |
| ARI_P_01 | 0.422 | -1.955 | 3.584 | 0.815 | 0.05 | 0.17 | 0.42 | 0.58 | 0.58 | 0.49 | 0.24 |
| ARI_P_02 | 2.089 | -0.507 | 1.681 | 0.587 | 0.01 | 0.08 | 0.8 | 1.49 | 0.3 | 0.82 | 1.48 |
| ARI_P_03 | 2.416 | 0.765 | 1.684 | 1.225 | 0.01 | 0.03 | 0.14 | 0.43 | 0.74 | 0.77 | 0.55 |
| ARI_P_04 | 4.090 | 1.304 | 2.139 | 1.722 | 0 | 0.01 | 0.05 | 0.17 | 0.42 | 0.65 | 0.65 |
| ARI_P_05 | 3.700 | 0.400 | 1.535 | 0.968 | 0 | 0 | 0.01 | 0.31 | 2.4 | 0.5 | 0.15 |
| ARI_P_06 | 2.251 | -0.017 | 1.642 | 0.813 | 0 | 0.04 | 0.33 | 1.33 | 0.68 | 0.8 | 1.22 |
| ARI_P_07 | 2.254 | 0.138 | 1.319 | 0.729 | 0.01 | 0.04 | 0.24 | 0.89 | 0.91 | 1.01 | 0.67 |
| Mean | 2.460 | 0.018 | 1.941 | 0.979 |  |  |  |  |  |  |  |
| Test Info |  |  |  |  | 0.08 | 0.37 | 1.99 | 5.19 | 6.03 | 5.05 | 4.97 |
| SEM |  |  |  |  | 3.55 | 1.64 | 0.71 | 0.44 | 0.41 | 0.45 | 0.45 |
| Reliability |  |  |  |  | -11.62 | -1.69 | 0.5 | 0.81 | 0.83 | 0.8 | 0.8 |

**Supplementary Table 7.** *Item Response Theory Parameters for MFQ*.

| | Discrimination ($\alpha$) and difficulty ($\beta$) parameters | | | | Item information by area of the latent trait (z-score) | | | | | | |
| --- | --- | --- | --- | --- | --- | --- | --- | --- | --- | --- | --- |
| | $\alpha$ | $\beta_1$ | $\beta_2$ | Mean($\beta$) | -3 | -2 | -1 | 0 | 1 | 2 | 3 |
| MFQ_P_01 | 2.085 | 0.137 | 1.868 | 1.003 | 0.03 | 0.11 | 0.3 | 0.5 | 0.54 | 0.52 | 0.33 |
| MFQ_P_02 | 1.902 | 1.271 | 2.896 | 2.084 | 0.01 | 0.03 | 0.1 | 0.24 | 0.43 | 0.52 | 0.47 |
| MFQ_P_03 | 1.148 | 1.579 | 3.487 | 2.533 | 0.04 | 0.07 | 0.12 | 0.17 | 0.2 | 0.2 | 0.17 |
| MFQ_P_04 | 0.776 | 1.507 | 3.904 | 2.706 | 0.05 | 0.07 | 0.09 | 0.11 | 0.12 | 0.11 | 0.1 |
| MFQ_P_05 | 1.257 | 1.369 | 2.733 | 2.051 | 0.04 | 0.08 | 0.14 | 0.23 | 0.29 | 0.28 | 0.21 |
| MFQ_P_06 | 1.489 | 1.731 | 3.290 | 2.511 | 0.02 | 0.05 | 0.11 | 0.19 | 0.28 | 0.32 | 0.28 |
| MFQ_P_07 | 1.342 | 0.820 | 2.493 | 1.657 | 0.05 | 0.09 | 0.16 | 0.24 | 0.28 | 0.26 | 0.19 |
| MFQ_P_08 | 2.738 | 1.206 | 2.557 | 1.882 | 0 | 0.01 | 0.06 | 0.2 | 0.49 | 0.63 | 0.63 |
| MFQ_P_09 | 1.710 | 1.379 | 2.876 | 2.128 | 0.02 | 0.05 | 0.11 | 0.24 | 0.39 | 0.45 | 0.37 |
| MFQ_P_10 | 1.302 | 0.431 | 2.308 | 1.370 | 0.05 | 0.11 | 0.19 | 0.28 | 0.31 | 0.27 | 0.19 |
| MFQ_P_11 | 1.442 | -0.401 | 1.747 | 0.673 | 0.08 | 0.16 | 0.27 | 0.34 | 0.34 | 0.26 | 0.16 |
| MFQ_P_12 | 1.503 | 1.130 | 2.914 | 2.022 | 0.02 | 0.06 | 0.14 | 0.27 | 0.4 | 0.43 | 0.34 |
| MFQ_P_13 | 1.510 | 2.574 | 3.859 | 3.217 | 0.02 | 0.03 | 0.07 | 0.14 | 0.23 | 0.31 | 0.32 |
| MFQ_P_14 | 1.355 | 1.273 | 2.801 | 2.037 | 0.03 | 0.07 | 0.14 | 0.24 | 0.32 | 0.32 | 0.24 |
| MFQ_P_15 | 2.515 | 1.499 | 2.487 | 1.993 | 0 | 0.01 | 0.05 | 0.16 | 0.42 | 0.68 | 0.67 |
| MFQ_P_16 | 2.615 | 1.810 | 2.906 | 2.358 | 0 | 0.01 | 0.03 | 0.11 | 0.29 | 0.53 | 0.62 |
| MFQ_P_17 | 1.903 | 1.928 | 3.131 | 2.530 | 0.01 | 0.03 | 0.07 | 0.16 | 0.3 | 0.41 | 0.4 |
| MFQ_P_18 | 2.519 | 1.897 | 3.486 | 2.692 | 0.01 | 0.01 | 0.04 | 0.11 | 0.25 | 0.39 | 0.44 |
| MFQ_P_19 | 2.388 | 2.123 | 3.638 | 2.881 | 0.01 | 0.02 | 0.04 | 0.1 | 0.22 | 0.36 | 0.43 |
| MFQ_P_20 | 1.770 | 2.039 | 3.478 | 2.759 | 0.01 | 0.03 | 0.08 | 0.15 | 0.27 | 0.35 | 0.35 |
| MFQ_P_21 | 1.188 | 0.460 | 2.377 | 1.419 | 0.06 | 0.11 | 0.17 | 0.23 | 0.25 | 0.22 | 0.16 |
| MFQ_P_22 | 1.786 | 1.539 | 3.055 | 2.297 | 0.01 | 0.04 | 0.09 | 0.21 | 0.36 | 0.44 | 0.4 |
| MFQ_P_23 | 3.357 | 1.462 | 2.537 | 2.000 | 0 | 0 | 0.02 | 0.08 | 0.32 | 0.72 | 0.74 |
| MFQ_P_24 | 2.294 | 1.398 | 2.849 | 2.124 | 0.01 | 0.03 | 0.08 | 0.2 | 0.38 | 0.5 | 0.49 |
| MFQ_P_25 | 1.799 | 1.606 | 2.921 | 2.264 | 0.01 | 0.03 | 0.09 | 0.2 | 0.38 | 0.49 | 0.44 |
| MFQ_P_26 | 1.101 | 1.817 | 3.531 | 2.674 | 0.04 | 0.07 | 0.1 | 0.14 | 0.16 | 0.17 | 0.14 |
| MFQ_P_27 | 1.472 | 1.303 | 3.132 | 2.218 | 0.03 | 0.06 | 0.13 | 0.23 | 0.33 | 0.36 | 0.29 |
| MFQ_P_28 | 2.098 | 1.454 | 2.897 | 2.176 | 0.01 | 0.03 | 0.08 | 0.2 | 0.39 | 0.52 | 0.5 |
| MFQ_P_29 | 1.554 | 0.716 | 2.363 | 1.540 | 0.03 | 0.08 | 0.18 | 0.32 | 0.41 | 0.39 | 0.27 |
| MFQ_P_30 | 1.755 | 0.969 | 2.499 | 1.734 | 0.02 | 0.06 | 0.15 | 0.3 | 0.44 | 0.46 | 0.34 |
| MFQ_P_31 | 2.205 | 1.112 | 3.118 | 2.115 | 0.01 | 0.03 | 0.1 | 0.24 | 0.41 | 0.44 | 0.44 |
| MFQ_P_32 | 1.118 | 1.377 | 3.886 | 2.632 | 0.04 | 0.07 | 0.12 | 0.16 | 0.2 | 0.2 | 0.17 |
| MFQ_P_33 | 1.066 | 1.834 | 3.713 | 2.774 | 0.04 | 0.07 | 0.1 | 0.14 | 0.17 | 0.17 | 0.15 |
| MFQ_P_34 | 2.245 | 1.384 | 2.548 | 1.966 | 0 | 0.02 | 0.06 | 0.2 | 0.47 | 0.66 | 0.62 |
| Mean | 1.774 | 1.345 | 2.950 | 2.147 |  |  |  |  |  |  |  |
| Test Info |  |  |  |  | 0.85 | 1.82 | 3.78 | 7.05 | 11.05 | 13.35 | 12.04 |
| SEM |  |  |  |  | 1.09 | 0.74 | 0.51 | 0.38 | 0.3 | 0.27 | 0.29 |
| Reliability |  |  |  |  | -0.18 | 0.45 | 0.74 | 0.86 | 0.91 | 0.93 | 0.92 |

**Supplementary Table 8.**
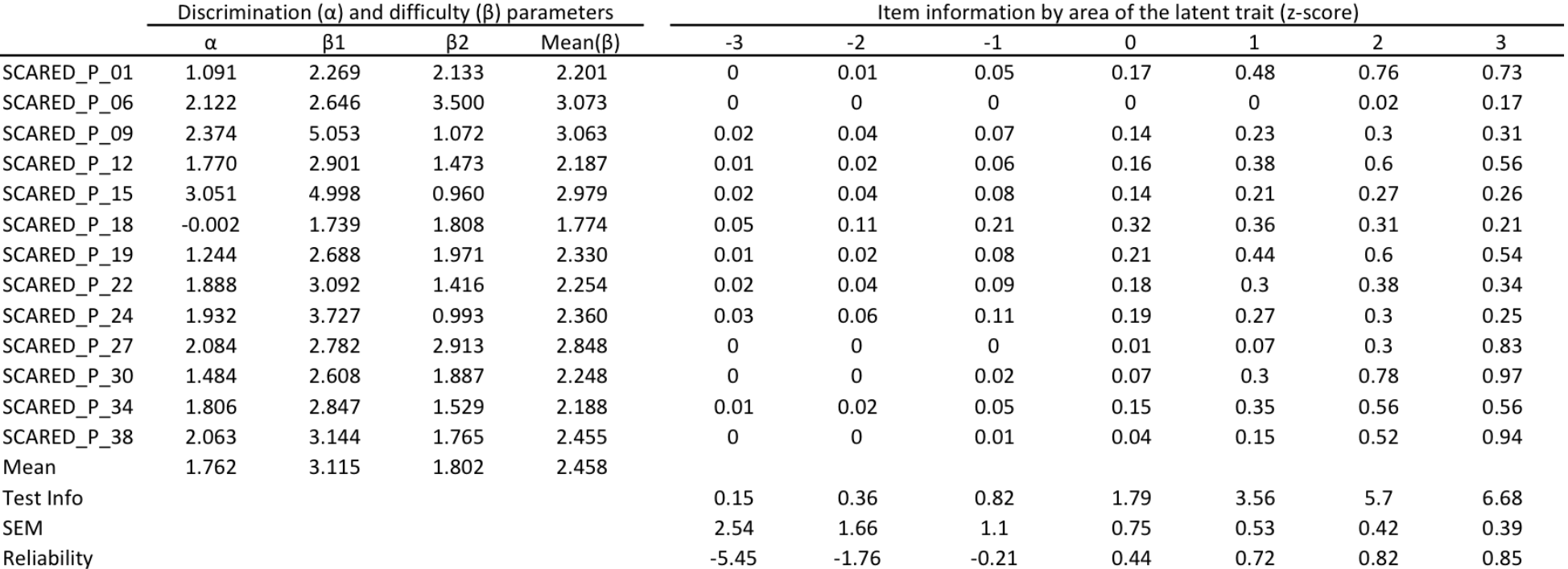
*Item Response Theory Parameters for SCARED Panic Disorder*.

**Supplementary Table 9.** *Item Response Theory Parameters for SCARED Social Anxiety*.

| | Discrimination ($\alpha$) and difficulty ($\beta$) parameters | | | | Item information by area of the latent trait (z-score) | | | | | | |
| --- | --- | --- | --- | --- | --- | --- | --- | --- | --- | --- | --- |
| | $\alpha$ | $\beta_1$ | $\beta_2$ | Mean($\beta$) | -3 | -2 | -1 | 0 | 1 | 2 | 3 |
| SCARED_P_03 | -0.198 | 1.399 | 1.874 | 1.637 | 0.06 | 0.15 | 0.32 | 0.47 | 0.49 | 0.38 | 0.2 |
| SCARED_P_10 | 0.268 | 1.807 | 2.165 | 1.986 | 0.02 | 0.07 | 0.22 | 0.47 | 0.58 | 0.57 | 0.39 |
| SCARED_P_26 | 0.248 | 1.274 | 3.533 | 2.404 | 0 | 0.02 | 0.13 | 0.82 | 1.11 | 0.83 | 1.19 |
| SCARED_P_32 | -0.189 | 0.998 | 4.152 | 2.575 | 0.01 | 0.11 | 0.94 | 1.24 | 0.31 | 1.07 | 1.11 |
| SCARED_P_39 | 0.365 | 1.454 | 1.596 | 1.525 | 0.05 | 0.11 | 0.23 | 0.36 | 0.41 | 0.33 | 0.19 |
| SCARED_P_40 | 0.284 | 1.598 | 1.847 | 1.723 | 0.04 | 0.1 | 0.23 | 0.39 | 0.47 | 0.39 | 0.23 |
| SCARED_P_41 | -0.026 | 1.466 | 2.517 | 1.992 | 0.03 | 0.1 | 0.34 | 0.62 | 0.63 | 0.64 | 0.39 |
| Mean | 0.107 | 1.428 | 2.526 | 1.977 |  |  |  |  |  |  |  |
| Test Info |  |  |  |  | 0.21 | 0.67 | 2.41 | 4.38 | 4.01 | 4.21 | 3.71 |
| SEM |  |  |  |  | 2.21 | 1.22 | 0.64 | 0.48 | 0.5 | 0.49 | 0.52 |
| Reliability |  |  |  |  | -3.87 | -0.49 | 0.59 | 0.77 | 0.75 | 0.76 | 0.73 |

